# Granulocyte colony-stimulating factor acts through calcium-permeable AMPA receptors to potentiate cocaine reward

**DOI:** 10.64898/2026.01.30.702629

**Authors:** Rebecca S. Hofford, Rashaun Wilson, Colin McArdle, Tanner Euston, Jonathon P. Sens, Abigail Mastrantoni, Taylor McCabe, Katherine R. Meckel, Weiwei Wang, Kelsey E. Lucerne, Aya Osman, Kimberly F. Raab-Graham, TuKiet T. Lam, Drew D. Kiraly

## Abstract

Neuroimmune interactions have emerged as critical modulators of substance use disorders and may represent promising translational therapeutic targets. In prior work, we demonstrated that the cytokine granulocyte colony stimulating factor (G-CSF) is elevated in mice following cocaine exposure, with circulating levels correlating with cocaine intake and locomotor sensitization. Additionally, exogenous G-CSF enhances cocaine reward and increases low-dose cocaine self-administration. We have further shown that repeated G-CSF administration alters expression of glutamatergic synapse-associated proteins following cocaine-seeking behavior. Building on these findings, the present studies examined the molecular consequences of repeated administration of G-CSF, cocaine, or their combination, with a focus on glutamatergic signaling pathways. We also tested whether altered glutamate receptor expression contributes to G-CSF-mediated enhancement of cocaine reward. Repeated combined administration of G-CSF and cocaine produced robust changes in glutamate-associated and synapse-related protein expression within the nucleus accumbens and medial prefrontal cortex. These molecular adaptations were accompanied by increased synaptic density in the nucleus accumbens. Finally, pharmacological inhibition of calcium-permeable AMPA receptors within the nucleus accumbens reversed the G-CSF-induced enhancement of cocaine conditioned place preference. Together, these findings indicate that G-CSF enhances cocaine reward at least in part by promoting glutamatergic synaptic remodeling in the nucleus accumbens, identifying a neuroimmune-glutamate mechanism that may be leveraged for therapeutic intervention.

## Introduction

Cocaine use disorder is a debilitating disease affecting millions in the United States yearly. It is defined by repeated cycles of active drug taking, abstinence, and relapse, which makes long term recovery extremely difficult. Given that there are no FDA approved treatments for this disorder and the consistently high rates of stimulant overdose^1^, the need for new targets and treatment approaches has never been more urgent. Recent evidence suggests the immune system might impact the development and maintenance of neuropsychiatric conditions such as anxiety, depression, and substance use disorders^2–4^, resulting in a new wave of pharmaceuticals focused on immune system modulation being developed to treat these diseases^5^.

Our lab has described a role for granulocyte colony-stimulating factor (G-CSF) in the behavioral and cellular effects of cocaine reward. G-CSF is a pleiotropic cytokine and growth factor named for its ability to stimulate production of leukocytes from bone marrow^6^. In our lab, we observed that levels of G-CSF were increased in serum after mice were given or self-administered cocaine, and levels of G-CSF correlated with cocaine locomotor sensitization and intake^7^. Exogenously administered G-CSF enhanced cocaine conditioned place preference (CPP) and cocaine self-administration at low doses^7^. Finally, the cocaine-induced increase in G-CSF was necessary for reward, as blockade of G-CSF signaling in nucleus accumbens (NAc) abolished cocaine CPP^7^.

Based on prior evidence that G-CSF improves behavioral flexibility^8^, improves cognition in models of Alzheimer’s disease^9–11^, and improves motor performance after stroke^12^, it is clear that G-CSF exerts pleiotropic effects on a multitude of brain circuits, but full understanding of its mechanism underlying cocaine reward remains uncertain. Work from our lab has shown that G-CSF acts directly in the NAc to enhance both basal and cocaine-induced dopamine release^8,13,14^. However, in another study, we found that glutamatergic systems might also contribute to the effects of G-CSF on cocaine reinstatement. Using an unbiased proteomic approach, we have shown that daily G-CSF administration during abstinence causes a robust decrease in proteins involved in glutamate transmission in medial prefrontal cortex (mPFC)^15^. This is in line with existing data from other fields; in disease models, G-CSF is protective against stroke-induced excitotoxic cell death by causing downregulation of glutamate transporters and receptors^12,16,17^.

Dopaminergic and glutamatergic neurotransmission converge within NAc to convey relevant information about drug-associated cues and rewarding stimuli^18,19^, so changes in signaling at either dopamine or glutamate synapses could have major effects on drug reward and reinforcement. However, given our prior proteomics data that identified differences in the expression of glutamate synaptic proteins and data suggesting G-CSF improves cognition in Alzheimer’s disease models^9,10^ and recovery of function after stroke by regulating glutamate release or binding sites^20,21^, this study tests the hypothesis that G-CSF affects cocaine reward in part by regulating glutamatergic signaling.

In the current study, we used unbiased proteomics together with Western blotting and proximity ligation assays to define how cocaine alone or in combination with G-CSF alters glutamate receptor expression and glutamatergic synapse density. These approaches were used to identify molecular and synaptic adaptations associated with G-CSF modulation of cocaine exposure. We then examined the functional relevance of these changes with a final mechanistic experiment testing whether calcium-permeable AMPA receptors within the nucleus accumbens are required for G-CSF-induced enhancement of cocaine reward.

## Materials and Methods

### Animals

Male C57BL6/J (Jackson Labs, 7-9 weeks old at the start of the experiment) were housed four or five per cage upon arrival to the colony. All mice were housed in a temperature and humidity-controlled vivarium on a 12hr light:dark cycle. All behavioral experiments and injections were performed during animals’ light cycle. All animal procedures were approved by the animal care committees of Mount Sinai School of Medicine or Wake Forest School of Medicine, and all animal procedures conformed to ARRIVE guidelines.

### Drug Preparations and Doses

Doses of G-CSF, cocaine, and G-CSF+cocaine were the same for every experiment. Cocaine hydrochloride was provided by the drug supply program at NIDA and was dissolved in 0.9% saline (for CPP experiments) or 1xPBS (for cellular and biochemical analyses) to a final dose of 7.5 mg/kg. Mouse G-CSF (GenScript, Piscataway NJ) was dissolved in 1xPBS to a final dose of 50 ug/kg. Combination G-CSF+cocaine was given as a single injection and consisted of both G-CSF and cocaine dissolved in 1xPBS. All systemically injected solutions were injected at a volume of 10ml/kg. 1-naphthylacetyl spermine (NASPM, Tocris Bioscience) was dissolved in saline.

### Injections for cellular and biochemical assays

All mice were assigned to one of four groups: PBS, G-CSF, cocaine, or G-CSF+cocaine. Mice received their injections (i.p.) once daily for 7 days. Tissue was collected 24 hours after their last injection.

### Data-Independent Acquisition (DIA)

Mice were euthanized and whole tissue punches from the NAc (n=5-8/group) and mPFC (n=7-8/group) were acquired and flash frozen. Brain regions were identified using a brain matrix-the anterior commissure was used as a landmark for NAc and the corpus callosum as a landmark for mPFC. DIA LC–MS/MS was performed using a nanoACQUITY UPLC system (Waters Corporation, Milford, MA, USA) connected to a Q-Exactive HFX (ThermoFisher Scientific, San Jose, CA, USA) mass spectrometer. After injection, the samples were loaded into a trapping column (nanoEase M/Z Symmetry C18 Trap column, 180 µm × 20 mm) for 3 minutes at a flow rate of 10 µL/min and separated with a C18 column (nanoEase M/Z column Peptide BEH C18, 75 µm × 250 mm). The compositions of mobile phases A and B were 0.1% formic acid in water and 0.1% formic acid in ACN, respectively. The peptides were separated and eluted with a gradient extending from 6% to 25% mobile phase B in 98 min and then to 85% mobile phase B in additional 5 min at a flow rate of 300 nL/min and a column temperature of 37 °C. Column regeneration and up to three blank injections were carried out in between all sample injections. The data were acquired with the Q-Exactive HFX mass spectrometer operating in a data-independent acquisition mode with an isolation window width of 10 m/z. The full scan was performed in the range of 400–1,000 m/z with “Use Quadrupole Isolation” enabled at an Orbitrap resolution of 120,000 at 200 m/z and automatic gain control (AGC) target value of 3 × 10^6^. Fragment ions from each peptide MS^2^ were generated in the C-trap with higher-energy collision dissociation (HCD) at a normalized collision energy of 28% and detected in the Orbitrap at a resolution of 30,000.

DIA spectra were searched against a *Mus Musculus* brain proteome fractionated spectral library generated from DDA LC MS/MS spectra (collected from the same Q-Exactive HFX mass spectrometer) using Scaffold DIA software v. 2.2.0 (Proteome Software, Portland, OR, USA). Within Scaffold DIA, raw files were first converted to the mzML format using ProteoWizard v. 3.0.11748. The samples were then aligned by retention time and individually searched with a mass tolerance of 10 ppm and a fragment mass tolerance of 10 ppm. The data acquisition type was set to “Non-Overlapping DIA”, and the maximum missed cleavages was set to 2. Fixed modifications included carbamidomethylation of cysteine residues (+57.02). Dynamic modifications included phosphorylation of serine, threonine, and tyrosine (+79.96), deamination of asparagine and glutamine (+0.98), oxidation of methionine and proline (+15.99), and acetylation of lysine (+42.01). Peptides with charge states between 2 and 4 and 6–30 amino acids in length were considered for quantitation, and the resulting peptides were filtered by Percolator v. 3.01 at a threshold FDR of 0.01. Peptide quantification was performed by EncyclopeDIA v. 0.6.12 and six of the highest quality fragment ions were selected for quantitation. Proteins containing redundant peptides were grouped to satisfy the principles of parsimony, and proteins were filtered at a threshold of two peptides per protein and an FDR of 1%. The mass spectrometry proteomics data have been deposited to the ProteomeXchange Consortium via the PRIDE^22^ partner repository with the dataset identifier PXD070033.

### Pathway analysis and visualization of protein networks

Proteins were excluded from analysis if they were not detected in > 50% of all samples irrespective of treatment. Pairwise comparisons of the Log_10_ median intensity of every remaining protein were made using Scaffold DIA proteomics analysis software. Biological replicates were treated as independent samples and proteins were considered significantly differentially regulated when *p*<0.05. All groups were compared to saline to allow for inferences across comparisons. Protein names were converted to gene names prior to pathway analysis. The list of significantly differentially regulated proteins were uploaded into the open source pathway analysis software package G:Profiler^23^ to identify significantly enriched Gene Ontologies (GO) using an FDR corrected *p*<0.05. A list of shared proteins that were regulated in the same direction between treatment groups was generated using BioVenn^24^; non-overlapping proteins differentially regulated in cocaine treated and G-CSF+cocaine groups were separately uploaded into the STRING database^25^. Significant pathways (FDR corrected *p*<0.05) were sorted and displayed using pathway strength [log_10_(observed/expected)]. Unique proteins in GO pathway names containing the word “synapse” were counted for each comparison using G:Profiler. Portions of figures 1 and 2 were made with Biorender.com with full permission to publish.

**Fig. 1.**
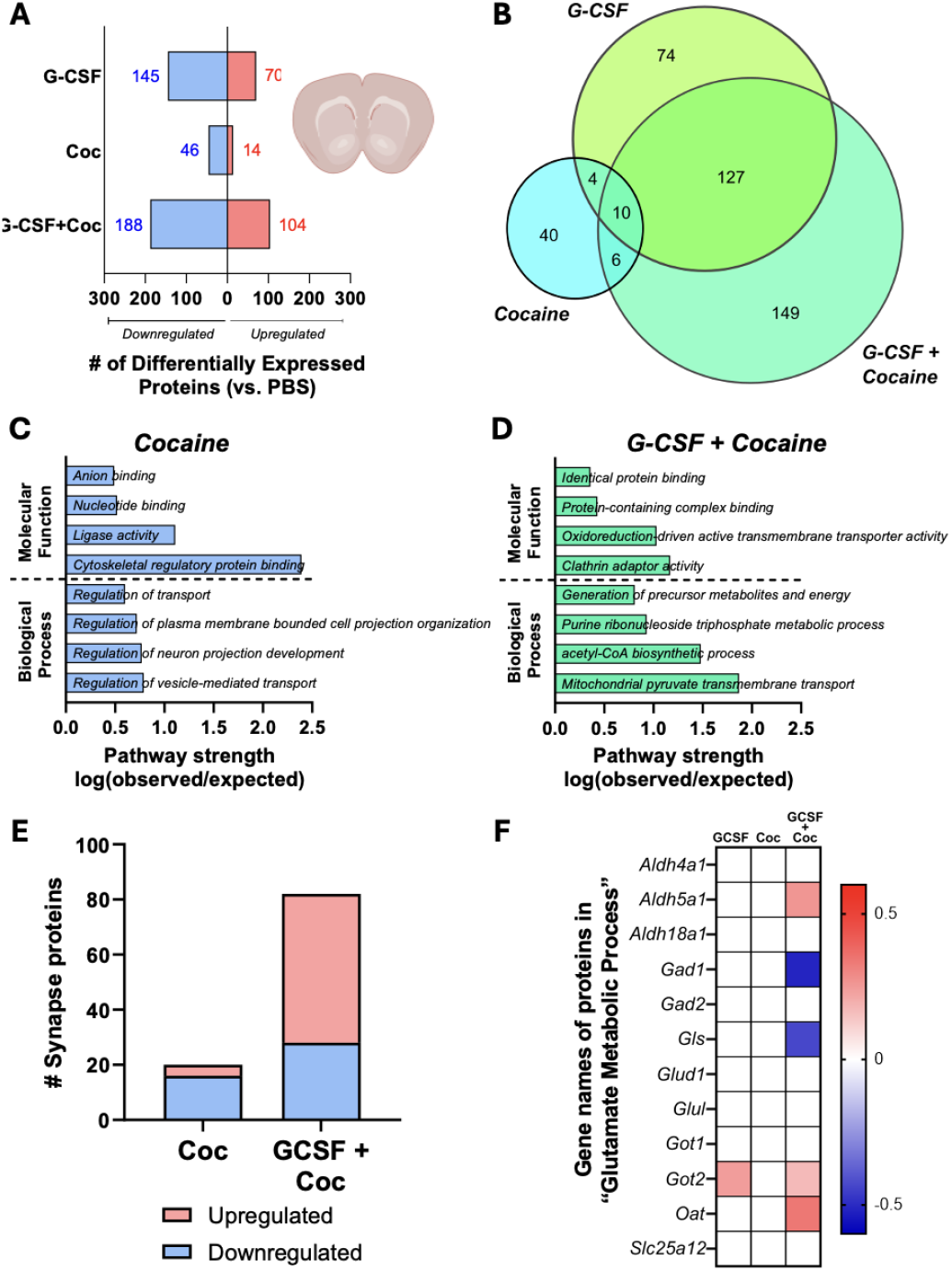
G-CSF + cocaine combination treatment alters expression of synapse- and glutamate-related proteins in nucleus accumbens. A. The number of differentially expressed proteins in mice treated with G-CSF (50 ug/kg), cocaine (7.5 mg/kg), or G-CSF+cocaine (50 ug/kg + 7.5 mg/kg) compared to saline. Downregulated proteins are indicated by blue bars and upregulated by red bars. B. Venn diagram of differentially expressed proteins from mice treated with G-CSF, cocaine, or G-CSF+cocaine compared to saline. C. Top significant biological process and molecular function GO pathways from cocaine treated mice by pathway strength. D. Top significant biological process and molecular function GO pathways from G-CSF+cocaine treated mice by pathway strength. E. The number of differentially regulated proteins in synapse-related pathways from cocaine and G-CSF+cocaine groups. Upregulated proteins are red and downregulated are blue. F. Heatmap of differentially regulated proteins in the “glutamate metabolic process” pathway in all treatment groups compared to saline. Color indicates fold change with blue as downregulated and red as upregulated. Proteins without a color indicator were not significantly differentially regulated. Only proteins detected in NAc are shown.

**Fig. 2.**
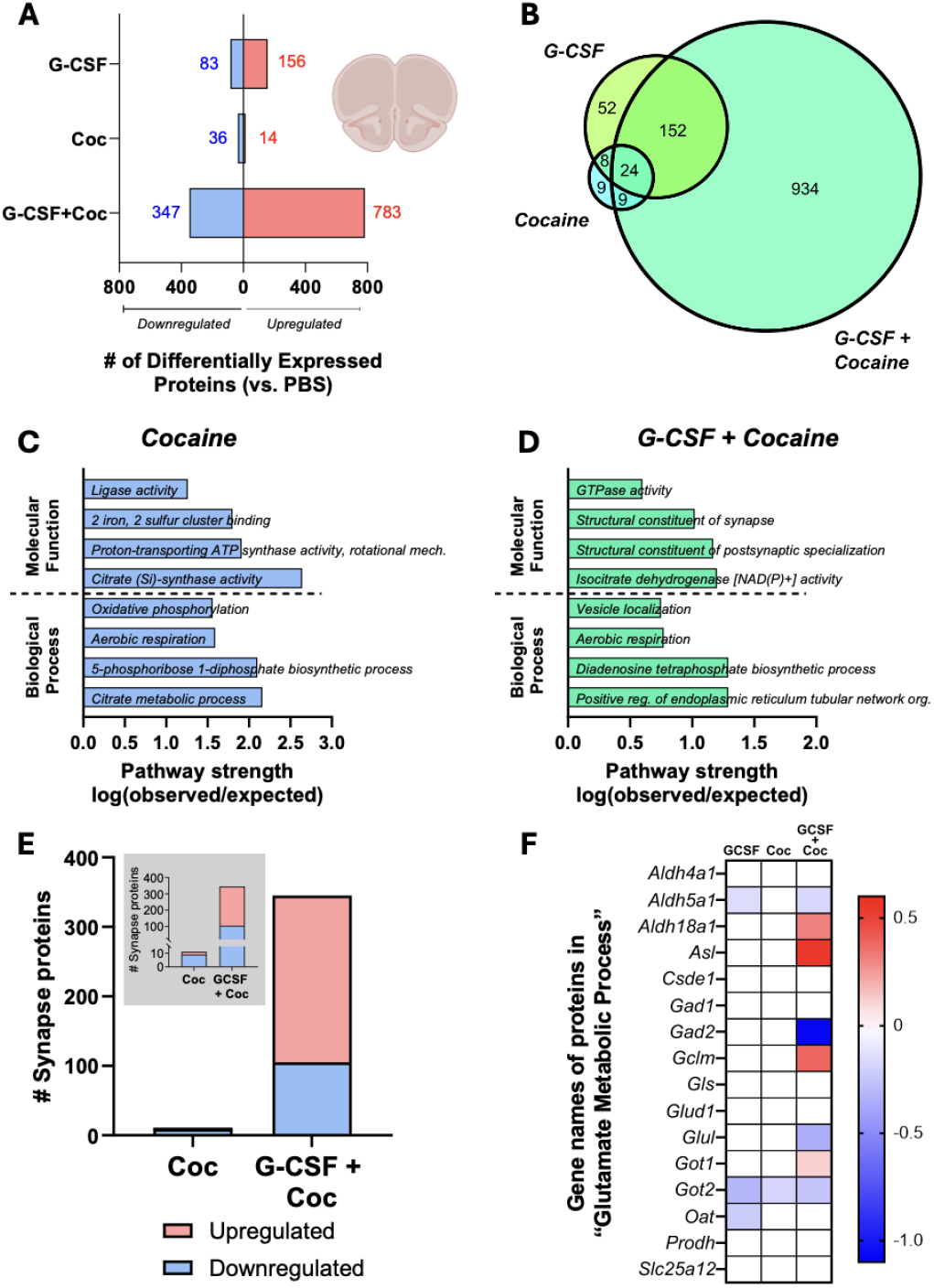
G-CSF + cocaine combination treatment alters expression of synapse- and glutamate-related proteins in medial prefrontal cortex. A. The number of differentially expressed proteins in mice treated with G-CSF (50 ug/kg), cocaine (7.5 mg/kg), or G-CSF+cocaine (50 ug/kg + 7.5 mg/kg) compared to saline. Downregulated proteins are indicated by blue bars and upregulated by red bars. **B**. Venn diagram of differentially expressed proteins from mice treated with G-CSF, cocaine, or G-CSF+cocaine compared to saline. **C**. Top significant biological process and molecular function GO pathways from cocaine treated mice by pathway strength. **D**. Top significant biological process and molecular function GO pathways from G-CSF+cocaine treated mice by pathway strength. **E**. The number of differentially regulated proteins in synapse-related pathways from cocaine and G-CSF+cocaine groups. Upregulated proteins are red and downregulated are blue. **F**. Heatmap of differentially regulated proteins in the “glutamate metabolic process” pathway in all treatment groups compared to saline. Color indicates fold change with blue as downregulated and red as upregulated. Proteins without a color indicator were not significantly differentially regulated. Only proteins detected in mPFC are shown.

### Proximity ligation assay

Twenty-four hours after their last injection, mice were transcardially perfused with EDTA-1xPBS followed by 4% paraformaldehyde. Brains were cryoprotected in 30% sucrose before freezing and brains were sectioned at 25um. Sections containing the NAc and mPFC were collected from each sample (n = 3 mice/gp; 1-3 slices/mouse). Sections were first blocked (10% normal donkey serum, 0.25% Tween-20) for two hours, followed by an overnight incubation with antibodies for chicken anti-MAP2 (1:500, Abcam ab5392), anti-PSD95 (1:400, Neuromab 75-028), and anti-synapsin1 (1:200, Santa Cruz s.c.7379). After washing, sections were incubated for two hours at 37ºC in secondary antibodies (anti-mouse PLUS (1:5, Sigma-Aldrich DUO92001) and anti-goat MINUS (1:5, Sigma-Aldrich DUO92006) followed by sequential steps of ligation and amplification per assay instructions (Sigma-Aldrich, DUO92008). Confocal images were taken on a Nikon A1plus confocal microscope at 20X air magnification after focusing on clear MAP2 staining. Images were analyzed using the ImageJ (FIJI) “analyze particles” function after normalization of threshold across images of the same brain area. Number of particles in the region of interest (ROI) was normalized for size of the region by dividing the number of puncta detected by area of ROI. Fold change in puncta/area was calculated and graphed by dividing each slice’s normalized puncta count by the average normalized puncta count for control mice.

### Western blot

Mice were injected with PBS, G-CSF, cocaine, or combination G-CSF+cocaine, euthanized, and tissue punches were collected as described above (n=9-10/gp). Sections were homogenized in lysis buffer (5% Tris-HCl, 10% SDS, 10% NaCl, 1% 0.5M EDTA, and protease inhibitor (ThermoFisher Scientific)) using a sonicator. Total protein from the resulting lysates were calculated using the Pierce BCA protein assay (ThermoFisher Scientific). Ten ug of protein were added to each well of a 4-12% polyacrylamide gel (Invitrogen) alongside a protein ladder (Bio-Rad, Precision Plus Dual Color) and electrophoresis was run at 100V until dye ran off the end of the gel. Proteins were transferred to a PVDF membrane and blocked with Intercept blocking buffer (LiCor) before incubation with GluR1 (1:1000, Cell Signaling Technology #13185), PSD95 (1:1000, Cell Signaling Technology #36233), or GluR2 (1:1000, Cell Signaling Technology #13607), and actin (1:5000, MP Biomedicals #691001) antibodies overnight at 4°C. Blots were incubated with secondary antibodies IRDye800CW anti-rabbit and IRDye680RD anti-mouse before imaging on an Odyssey LiCor. Staining intensity was analyzed using the blot analysis function in ImageJ. Each sample’s target protein staining intensity (GluR1, GluR2, or PSD95) was normalized to their loading control staining intensity (actin) by dividing target intensity by actin intensity. The resulting proportions were normalized to control samples (mice receiving PBS) and converted to percent control.

### Implantation of bilateral cannula into NAc

Mice were anesthetized with ketamine/xylazine (10 mg/kg / 100 mg/kg, i.p.) before placement in a stereotaxic frame. Bilateral cannula (P1 Technologies) were placed above the NAc using the coordinates: A/P +1.6mm, M/L +/-1.0mm, and D/V + 4.0mm and were secured to the skull using dental acrylic and a skull screw anchor. Cannula caps and dummy cannulas were used to prevent debris from entering the cannula. Mice were allowed to recover for one week minimum.

### Conditioned place preference

Cocaine conditioned place preference was performed on mice (n=6-7/group) between one and two weeks after stereotaxic cannula implant using an unbiased method as described previously^7,26^. The CPP procedure occurred over four consecutive days. On pre-test day (day 1), mice received injections of PBS or G-CSF one hour before placement into the center chamber of the CPP apparatus. On conditioning days (days 2 and 3), all mice received injections of 1xPBS or G-CSF one hour before they were injected with saline. Three hours after saline conditioning session, mice received injections of 7.5 mg/kg cocaine (i.p.) before placement in their cocaine chamber. On test day (day 4), mice received PBS or G-CSF 45 mins before intra-NAc infusion of 0.5 ul of vehicle or 20 ug of the calcium permeable AMPA receptor antagonist NASPM/side over 5 minutes. Fifteen minutes after infusion end, mice were placed in the center chamber of the CPP apparatus and were allowed to explore all three chambers. Cocaine place preference was calculated as: [time in drug-paired chamber on test day – time in saline-paired chamber on test day]. To confirm cannula placement, 1 ul/side Evan’s blue dye (Sigma Aldrich) was infused bilaterally into NAc 24 hrs after CPP test. Mice were then euthanized and brains were flash frozen and sliced.

### Statistical analysis

For Western blot data, signal between treatment groups was analyzed using separate 2-way ANOVAs with cocaine treatment (vehicle or cocaine) and G-CSF treatment (PBS or G-CSF) as between-subjects factors. To determine differences in synaptic density via the proximity ligation assay, a 2-way ANOVA with cocaine treatment and G-CSF treatment as between-subjects factors was used. Significance for the conditioned place preference was assessed using a 2-way ANOVA with G-CSF treatment and NASPM dose as between-subjects factors. All primary significance tests were followed by *a priori* pairwise comparison using Fisher’s LSD multiple comparison test.

## Results

### Proteins involved in synaptic signaling and glutamate metabolism are significantly altered in the NAc and mPFC of mice receiving G-CSF+cocaine

Since we have previously observed robust effects of G-CSF on enhancement of low dose cocaine CPP and cocaine self-administration^7^, we wanted to investigate the proteomic landscape within the reward circuitry of mice receiving G-CSF, cocaine, or G-CSF+cocaine. We utilized data-independent acquisition to analyze global changes in protein expression in an unbiased manner. G-CSF alone altered the expression of 215 proteins within the NAc, a larger number of proteins than cocaine alone, which produced 60 differentially regulated proteins (**Fig. 1A**). However, G-CSF+cocaine combination treatment had the greatest effect on protein expression, by up- and downregulating expression of 292 proteins (**Fig. 1A**). Most proteins that were differentially regulated by G-CSF alone or in combination with cocaine overlapped (∼64% of proteins altered by G-CSF alone overlapped with G-CSF+cocaine and ∼47% of G-CSF+cocaine proteins overlapped with G-CSF alone). Cocaine treatment produced a largely unique protein expression signature, with only a third of differentially regulated proteins overlapping with G-CSF alone or G-CSF+cocaine (**Fig. 1B**).

Since we primarily sought to understand how G-CSF modulates cocaine reward, further analysis focused on proteins that were uniquely regulated by cocaine (40 proteins) or G-CSF+cocaine (149 proteins). Pathway analysis of unique cocaine-regulated proteins identified pathways related to “cytoskeletal regulatory protein binding” (**Fig. 1C**), while G-CSF+cocaine altered expression of proteins involved in “mitochondrial pyruvate transmembrane transport” among others (**Fig. 1D**).

In addition to our unbiased examination of global protein changes, we examined expression of glutamatergic synapse-related proteins since our prior work found that G-CSF caused robust downregulation of proteins involved in glutamatergic synaptic signaling after cocaine abstinence. To determine if any meaningful differences existed in the expression of synapse related proteins, we determined the number of proteins that were up- or downregulated in any “synapse”-related pathways. Cocaine treatment led to an upregulation of four and downregulation of 16 synapse-related proteins but G-CSF+cocaine treatment upregulated 54 and downregulated 28 (**Fig. 1E**), suggesting that G-CSF+cocaine treatment produced a larger percentage of upregulated to downregulated synapse related proteins while cocaine had a larger percentage of downregulated proteins.

Additionally, closer examination of all pathways related to “glutamate” noted an interesting glutamate-related pathway significant only in G-CSF+cocaine: “glutamate metabolic process” (GO:00065). This pathway contains 28 proteins total, 12 of which were detected in NAc. G-CSF+cocaine treatment altered the expression of five glutamate metabolism proteins including downregulation of glutamate decarboxylase 1 and glutaminase as well as upregulation of proteins encoded by *Aldh5a1, Got2*, and *Oat* (**Fig. 1F**). Only one glutamate metabolism protein was upregulated by G-CSF alone and none were altered by cocaine.

Within mPFC, G-CSF treatment also altered expression of more proteins (239) than cocaine alone (50). Similar to the NAc, G-CSF+cocaine produced the largest number of differentially regulated proteins in any comparison with 1,130 proteins (**Fig. 2A**), the vast majority of which did not overlap with G-CSF alone or cocaine alone (**Fig. 2B**). Unique proteins regulated by cocaine generated pathways included “citrate metabolic process” among others (**Fig. 2C**). These pathways differed from those predicted by G-CSF+cocaine and included “structural component of synapse” and others (**Fig. 2D**).

*A priori* investigation of synapse and glutamate related proteins generated similar findings to that observed in the NAc. Cocaine had two synapse-related proteins that were upregulated and nine proteins downregulated and G-CSF+cocaine upregulated 240 synapse-related proteins and downregulated 105 (**Fig. 2E**). Similarly, G-CSF+cocaine altered the expression of eight glutamate metabolism related proteins (of 16 total found in mPFC), while G-CSF alone downregulated three and cocaine downregulated one (**Fig. 2F**).

### G-CSF+cocaine increases glutamatergic synaptic density in NAc over cocaine alone

Since we observed increased overall expression of synaptic proteins in mice treated with G-CSF+cocaine, we next utilized a proximity ligation assay (PLA) to assess whether G-CSF+cocaine increases glutamatergic synapse number within the reward circuit. PLA fluorescently labels a set of proteins that are located within 40nm of each other. Since synapses are 40nm or smaller, PLA can be used to label functional synapses^27,28^. This method has advantages over other spine measuring techniques, in that synapses are labeled only if there is both a presynaptic and postsynaptic element present. We utilized synapsin 1 and PSD-95 to specifically label functional glutamatergic synapses^28,29^. Within NAc, we found that cocaine-treated mice had more glutamatergic synapses than mice treated with vehicle (**Fig. 3A**; significant effect of cocaine: F_(1, 26)_=12.43; *p*=0.002; significant interaction: F_(1,26)_=10.52, *p*=0.003; effect of G-CSF: F_(1,26)_=3.39, *p*=0.071). Additionally, pairwise comparisons indicated that G-CSF+cocaine treated mice had higher synaptic density than mice treated with cocaine alone (G-CSF effect in saline: *p*=0.31; G-CSF effect in cocaine: *p*=0.002). In mPFC, neither G-CSF nor cocaine affected synapse number within mPFC (**Fig. 3B**; effect of cocaine: F_(1,12)_=0.005; *p*=0.95; effect of G-CSF: F_(1,12)_=0.74, *p*=0.41; interaction: F_(1,12)_=1.43, *p*=0.26, no significant pairwise comparisons).

**Fig. 3.**
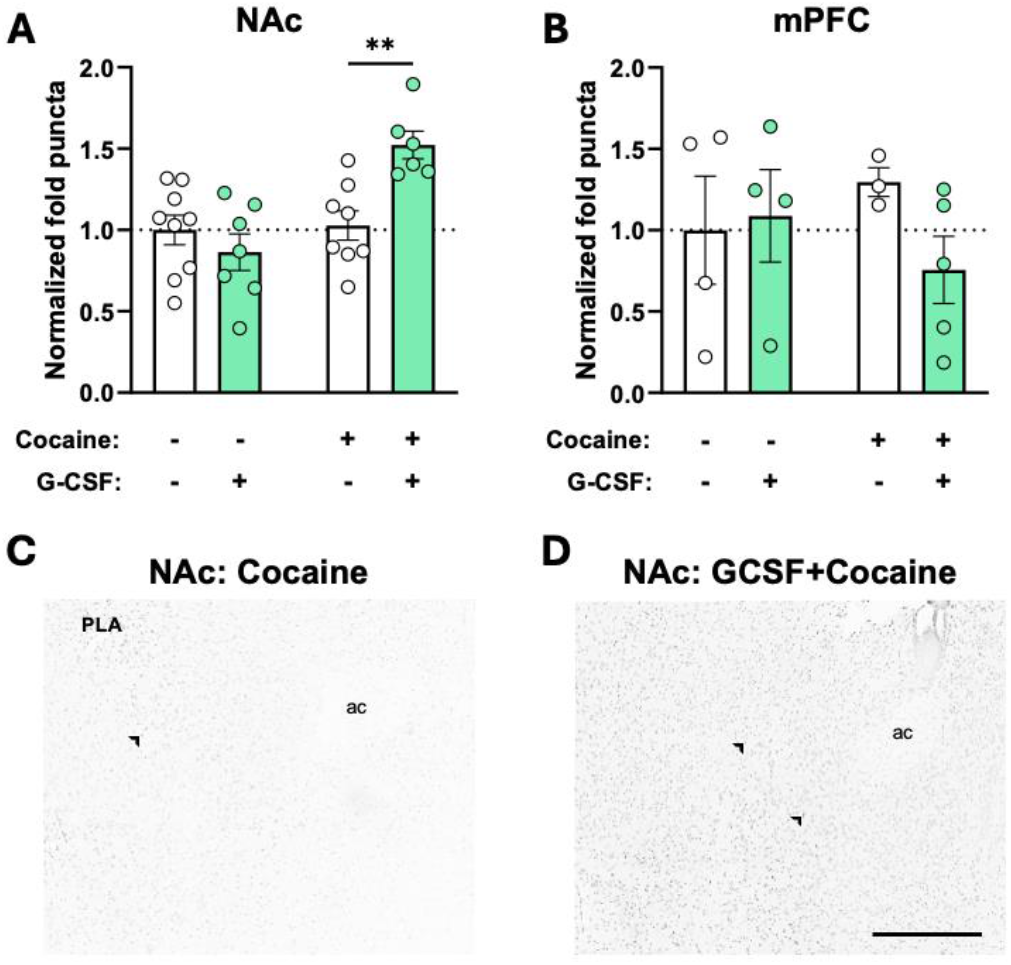
GCSF + cocaine increases glutamatergic synaptic density compared to cocaine alone in NAc but not mPFC. A. Fold change puncta intensity after normalization in NAc after treatment with saline, G-CSF, cocaine, or G-CSF+cocaine. B. Fold change puncta intensity after normalization in mPFC after treatment with saline, G-CSF, cocaine, or G-CSF+cocaine. C. Representative image of NAc from a cocaine treated mouse. D. Representative image of NAc from a G-CSF+cocaine treated mouse. Scale bar 500 μm. \*\**p <* 0.01. ac= anterior commissure.

### Expression of some glutamatergic synapse-associated proteins are upregulated after G-CSF+cocaine, but not cocaine alone

Given the current data that identified an increase in glutamatergic synaptic density within NAc (**Fig. 3**), we measured expression of the synaptic glutamatergic proteins, PSD95, GluR1, and GluR2 within NAc. For PSD95, there was a significant main effect of G-CSF only (**Fig. 4A**; G-CSF: F_(1,33)_=4.46, *p*=0.04; cocaine: F_(1,33)_=1.24, *p*=0.27; interaction: F_(1,33)_=0.91; *p*=0.35). *A priori* pairwise comparisons indicated that G-CSF+cocaine significantly increased PSD95 expression compared to PBS/vehicle (*p*=0.03) and PBS/cocaine (*p*=0.04). While there were no significant main effects on GluR1 expression (**Fig. 4B**; G-CSF: F_(1,32)_=0.22, *p*=0.15; cocaine: F_(1,32)_=2.25, *p*=0.14; interaction: F_(1,32)_=0.31, *p*=0.58), pairwise comparisons indicated that G-CSF+cocaine increased GluR1 expression compared to PBS/vehicle (*p*=0.04). Levels of GluR2 did not differ by treatment (**Fig. 4C**).

**Fig. 4.**
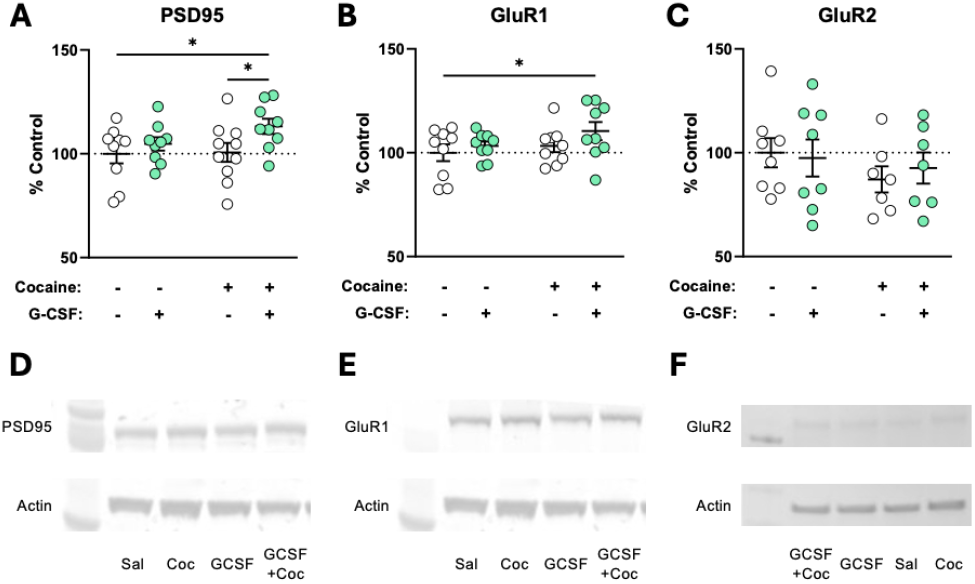
GCSF + cocaine increases expression of PSD95 and GluR1, not GluR2 within NAc. A.Percent change PSD95 expression from saline (control) in mice treated with G-CSF (50 ug/kg), cocaine (7.5 mg/kg), or G-CSF+cocaine (50 ug/kg + 7.5 mg/kg). **B**. Percent change GluR1 expression from saline (control) in mice treated with G-CSF, cocaine, or G-CSF+cocaine. **C**. Percent change GluR2 expression from saline (control) in mice treated with G-CSF, cocaine, or G-CSF+cocaine. **D**. Example PSD95 blot. **E**. Example GluR1 blot. **F**. Example GluR2 blot. * *p <* 0.05.

**Fig. 5.**
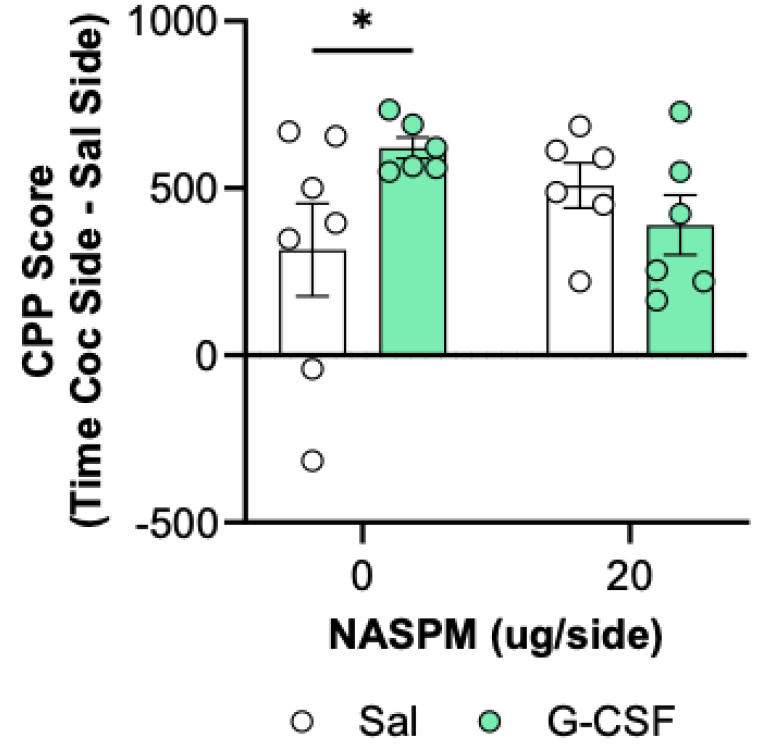
Intra-NAc NASPM reverses the effect of G-CSF on increased cocaine CPP. Mice given G-CSF display increased 7.5 mg/kg cocaine CPP when given saline into NAc. However, mice given 20 ug/side NASPM do not show enhanced cocaine CPP. *\* p <* 0.05.

### Antagonism of calcium-permeable AMPA receptors reverses the enhancement of cocaine CPP caused by G-CSF

Calcium-permeable AMPA receptors contain GluR1, but not GluR2 subunits; these AMPA receptors have been heavily implicated in cocaine craving after abstinence^30–32^. Since we observed an increase in GluR1, not GluR2, after G-CSF+cocaine, we hypothesized that calcium-permeable AMPA receptors in NAc might contribute to the effects of G-CSF on cocaine reward. Mice were treated daily with PBS or G-CSF during 7.5 mg/kg cocaine CPP. Before their CPP test, mice were given bilateral infusions of vehicle or 20ug of NASPM, an antagonist of calcium-permeable AMPA receptors into NAc. While G-CSF enhanced cocaine CPP in mice given vehicle (**Fig**. 5 - *p*=0.04), G-CSF had no effect on CPP in mice given 20ug of NASPM (*p*=0.56).

## Discussion

In the current set of studies, we found evidence that G-CSF, when given in combination with cocaine, increases the density of glutamatergic synapses within the NAc and increases expression of synapse-related proteins and proteins related to glutamate metabolism in both NAc and mPFC. We identified several proteins in the “glutamate metabolic process” pathway as being significantly differentially regulated in both NAc and mPFC of mice treated with combination G-CSF+cocaine. However, several enzymes are upregulated (*Aldh5a1, Got2*, and *Oat* in NAc and *Aldh18a1, Asl, Gclm*, and *Got1* in mPFC) while others are downregulated (*Gad1* and *Gls* in NAc and *Aldh5a1, Glul*, and *Got2* in mPFC). This pattern of up- and downregulation does not clearly indicate if G-CSF+cocaine treatment is promoting glutamate production or glutamate breakdown. It does, however, suggest that combination G-CSF+cocaine treatment affects glutamatergic signaling through two related mechanisms: alteration of metabolism directly and increases in synapse number and GluR1 expression.

In our previous work, we found that G-CSF modulated cocaine reinforcement and cocaine-seeking in opposite directions depending on when G-CSF was administered. If G-CSF was given during cocaine exposure, animals demonstrated increased place preference and self-administration of low dose cocaine^7^. However, when G-CSF was given during extinction or abstinence from cocaine self-administration, animals had faster extinction and reduced cocaine-seeking^15^. Similar to the behavioral results, G-CSF administration had opposite effects on the expression of glutamate synapse related proteins depending on when G-CSF was administered. In the current study, concurrent G-CSF and cocaine upregulated synapse-associated proteins and glutamate synapse number but when G-CSF was administered during abstinence from cocaine, we observed downregulation of glutamatergic receptors and associated proteins^15^. This raises the intriguing possibility that G-CSF bi-directionally modulates the effects of glutamate by either enhancing or reducing expression of glutamate-related proteins. G-CSF is known to reduce negative consequences of glutamate. In stroke models, G-CSF protects against excitotoxicity by downregulating glutamate receptors and transporters when extracellular glutamate is high^12,20,21^. However, more work needs to be done to understand how G-CSF works with cocaine to potentiate glutamate signaling.

Glutamate signaling within the NAc is crucial for the formation of drug-cue associations^33^. This is most evident during abstinence from cocaine self-administration, as the role of GluR1 on cocaine-seeking and incubation of craving has been extensively studied^30–32^. Surface expression of calcium-permeable AMPA receptors (i.e. those receptors that lack the GluR2 subunit and contain primarily GluR1 subunits) increases over time during abstinence in the NAc and blockade of these receptors reduces cocaine-seeking^30^. In the current study we see increased levels of total GluR1, not GluR2, 24 hours after administration of G-CSF+cocaine but not cocaine alone. This increase was behaviorally relevant; NASPM into the NAc abrogated the effect of G-CSF on enhanced cocaine CPP. Taken together, this suggests that G-CSF, when given concurrently with cocaine, accelerates molecular changes associated with cocaine abstinence by modulating glutamatergic signaling.

While this study indicates that G-CSF influences cocaine reward by increasing expression of GluR1 and enhancing signaling at glutamatergic synapses, more work needs to be done to synergize the current results with our prior work on G-CSF and dopamine release after cocaine^13,14^, since both glutamate and dopamine signaling within NAc are crucial for learning about drug-associated cues^33,34^. Additionally, while this study highlights the importance of G-CSF in modulating glutamatergic signaling, the specific cellular and molecular mechanisms underlying these changes remain unclear. G-CSF receptors in the brain are primarily expressed on microglia^35^ but some reports suggest direct neuron specific mechanisms of signaling^36,37^. Future studies will examine cellular and molecular mechanisms driving these changes. Regardless, the current work highlights how a cytokine, G-CSF, can modulate cocaine reward and provides evidence of its cellular mechanism. Crucially, this study supports the idea that immune modulators can be useful pharmacological tools for the treatment of mental health disorders.

## Data availability statement

Proteomics raw and analyzed data are available on ProteomeXChange as described in the Methods section. Raw and analyzed data from other experiments will be made available upon reasonable request.

## Acknowledgments

Cocaine hydrochloride was provided by the NIDA Drug Supply Program. We thank the Yale/NIDA Neuroproteomics Center (DA018343) for their support of these experiments. We also thank the Keck MS & Proteomics Resource at Yale School of Medicine for providing the mass spectrometers and the accompanying biotechnology tools for these studies, funded in part by the Yale School of Medicine and by the Office of The Director, National Institutes of Health (S10OD023651). The funders had no role in study design, data collection and analysis, decision to publish, or preparation of the manuscript.

## Author Contributions

RSH, RW, CM, TE, JPS, AM, TM, KRM, KEL, and AO performed experiments. TE, RSH, and DDK ran statistical analysis and made figures. DDK and RSH designed experiments. RSH wrote the manuscript with critical edits by KFRG, TTL, WW, KRM, and DDK. All authors approved the final manuscript.

## References

1. Garnett, M. F. & Minino, A. M. Drug Overdose Deaths in the United States, 2003–2023. (2024).

2. Hodes, G. E., Kana, V., Menard, C., Merad, M. & Russo, S. J. Neuroimmune mechanisms of depression. Nature Neuroscience 18, 1386–1393 (2015).

3. Hofford, R. S., Russo, S. J. & Kiraly, D. D. Neuroimmune mechanisms of psychostimulant and opioid use disorders. European Journal of Neuroscience 50, 2562–2573 (2019).

4. Lee, B. et al. Inflammatory and anti-inflammatory cytokines bidirectionally modulate amygdala circuits regulating anxiety. Cell 188, 2190-2202.e15 (2025).

5. Sharma, Y., Arora, M. & Bala, K. The potential of immunomodulators in shaping the future of healthcare. Discover Medicine 1, 37 (2024).

6. Lévesque, J.-P., Hendy, J., Winkler, I. G., Takamatsu, Y. & Simmons, P. J. Granulocyte colony-stimulating factor induces the release in the bone marrow of proteases that cleave c-KIT receptor (CD117) from the surface of hematopoietic progenitor cells. Experimental Hematology 31, 109–117 (2003).

7. Calipari, E. S. et al. Granulocyte-colony stimulating factor controls neural and behavioral plasticity in response to cocaine. Nature Communications 9, 9 (2018).

8. Kutlu, M. G. et al. Granulocyte Colony Stimulating Factor Enhances Reward Learning through Potentiation of Mesolimbic Dopamine System Function. J. Neurosci. 38, 8845 (2018).

9. Prakash, A., Medhi, B. & Chopra, K. Granulocyte colony stimulating factor (GCSF) improves memory and neurobehavior in an amyloid-β induced experimental model of Alzheimer’s disease. Pharmacology Biochemistry and Behavior 110, 46–57 (2013).

10. Tsai, K.-J., Tsai, Y.-C. & Shen, C.-K. J. G-CSF rescues the memory impairment of animal models of Alzheimer’s disease. Journal of Experimental Medicine 204, 1273–1280 (2007).

11. Sanchez-Ramos, J. et al. Granulocyte colony stimulating factor decreases brain amyloid burden and reverses cognitive impairment in Alzheimer’s mice. Neuroscience 163, 55–72 (2009).

12. Schäbitz, W.-R. et al. Neuroprotective Effect of Granulocyte Colony–Stimulating Factor After Focal Cerebral Ischemia. Stroke 34, 745–751 (2003).

13. Brady, L. J. et al. Granulocyte colony-stimulating factor (G-CSF) enhances cocaine effects in the nucleus accumbens via a dopamine release–based mechanism. Psychopharmacology 238, 3499–3509 (2021).

14. Brady, L. J., Hofford, R. S., Tat, J., Calipari, E. S. & Kiraly, D. D. Granulocyte-Colony Stimulating Factor Alters the Pharmacodynamic Properties of Cocaine in Female Mice. ACS Chem. Neurosci. 10, 4213–4220 (2019).

15. Hofford, R. S. et al. Granulocyte-Colony Stimulating Factor Reduces Cocaine-Seeking and Downregulates Glutamatergic Synaptic Proteins in Medial Prefrontal Cortex. J. Neurosci. 41, 1553 (2021).

16. Pan, C., Gupta, A., Prentice, H. & Wu, J.-Y. Protection of taurine and granulocyte colony-stimulating factor against excitotoxicity induced by glutamate in primary cortical neurons. Journal of Biomedical Science 17, S18 (2010).

17. Dumbuya, J. S., Chen, L., Wu, J.-Y. & Wang, B. The role of G-CSF neuroprotective effects in neonatal hypoxic-ischemic encephalopathy (HIE): current status. Journal of Neuroinflammation 18, 55 (2021).

18. Cooper, S., Robison, A. J. & Mazei-Robison, M. S. Reward Circuitry in Addiction. Neurotherapeutics 14, 687–697 (2017).

19. Vanderschuren, L. J. M. J. & Kalivas, P. W. Alterations in dopaminergic and glutamatergic transmission in the induction and expression of behavioral sensitization: a critical review of preclinical studies. Psychopharmacology 151, 99–120 (2000).

20. Han, J., Kollmar, R., Tobyas, B. & Schwab, S. Inhibited glutamate release by granulocyte-colony stimulating factor after experimental stroke. Neuroscience Letters 432, 167–169 (2008).

21. Mammele, S. et al. Prevention of an increase in cortical ligand binding to AMPA receptors may represent a novel mechanism of endogenous brain protection by G-CSF after ischemic stroke. Restorative Neurology and Neuroscience 34, 665–675 (2016).

22. Perez-Riverol, Y. et al. The PRIDE database resources in 2022: a hub for mass spectrometry-based proteomics evidences. Nucleic Acids Research 50, D543–D552 (2022).

23. Raudvere, U. et al. g:Profiler: a web server for functional enrichment analysis and conversions of gene lists (2019 update). Nucleic Acids Research 47, W191–W198 (2019).

24. Hulsen, T., de Vlieg, J. & Alkema, W. BioVenn – a web application for the comparison and visualization of biological lists using area-proportional Venn diagrams. BMC Genomics 9, 488 (2008).

25. Szklarczyk, D. et al. STRING v11: protein–protein association networks with increased coverage, supporting functional discovery in genome-wide experimental datasets. Nucleic Acids Research 47, D607–D613 (2019).

26. Hofford, R. S. et al. Alterations in microbiome composition and metabolic byproducts drive behavioral and transcriptional responses to morphine. Neuropsychopharmacol. 46, 2062–2072 (2021).

27. Heaney, C. F., McArdle, C. J. & Raab-Graham, K. F. DetectSyn: A Rapid, Unbiased Fluorescent Method to Detect Changes in Synapse Density. JoVE e63139 (2022) doi:10.3791/63139.

28. Heaney, C. F. et al. Role of FMRP in rapid antidepressant effects and synapse regulation. Molecular Psychiatry 26, 2350–2362 (2021).

29. El-Husseini, A., Schnell, E., Chetkovich, D., Nicoll, R. & Bredt, D. PSD-95 Involvement in Maturation of Excitatory Synapses. Science 290, 1364–1368.

30. Conrad, K. L. et al. Formation of accumbens GluR2-lacking AMPA receptors mediates incubation of cocaine craving. Nature 454, 118–121 (2008).

31. Lu, L., Grimm, J. W., Shaham, Y. & Hope, B. T. Molecular neuroadaptations in the accumbens and ventral tegmental area during the first 90 days of forced abstinence from cocaine self-administration in rats. Journal of Neurochemistry 85, 1604–1613 (2003).

32. Huang, Y. H. et al. In Vivo Cocaine Experience Generates Silent Synapses. Neuron 63, 40–47 (2009).

33. Britt, J. P. et al. Synaptic and Behavioral Profile of Multiple Glutamatergic Inputs to the Nucleus Accumbens. Neuron 76, 790–803 (2012).

34. Engel, L. et al. Dopamine neurons drive spatiotemporally heterogeneous striatal dopamine signals during learning. Current Biology 34, 3086-3101.e4 (2024).

35. Saunders, A. et al. Molecular Diversity and Specializations among the Cells of the Adult Mouse Brain. Cell 174, 1015-1030.e16 (2018).

36. Hasselblatt, M., Jeibmann, A., Riesmeier, B., Maintz, D. & Schäbitz, W.-R. Granulocyte-colony stimulating factor (G-CSF) and G-CSF receptor expression in human ischemic stroke. Acta Neuropathologica 113, 45–51 (2007).

37. Schneider, A. et al. The hematopoietic factor G-CSF is a neuronal ligand that counteracts programmed cell death and drives neurogenesis. J Clin Invest 115, 2083–2098 (2005).

